## Supplemental Material for "Potassium-mediated bacterial chemotactic response"

**The diffusion of potassium in semi-infinite liquid with a periodic source at $x = 0$.**

It has been found that the biofilm could lead a periodically oscillated source of potassium. Here, we simplified this source as a concentration signal of a cosine function at $x = 0$. Then the potassium will diffuse in the region of $x > 0$ which could be described by

$$\frac{\partial C}{\partial t} = D\frac{\partial^2 C}{\partial x^2}, \qquad \text{for } x > 0$$

with

$$C(x, t = 0) = 0,$$

$$C(x = 0, t) = \phi(t),$$

where $C$ is the concentration of potassium, $D$ is the diffusion constant of potassium in water, $L_0$ is the half maximum concentration of potassium, $T$ denotes the period of oscillation, and $\phi(t) = L_0(1 - \cos(2\pi t/T))$.

The solution can be given by[1]

$$C(x,t) = \int_0^t \phi(\lambda)\frac{\partial F(x, t-\lambda)}{\partial t}d\lambda,$$

where

$$F(x, t-\lambda) = \frac{2}{\sqrt{\pi}}\int_{\frac{x}{2\sqrt{D(t-\lambda)}}}^{\infty} e^{-\xi^2}d\xi$$

Thus

$$C(x,t) = \frac{2}{\sqrt{\pi}}\int_{\frac{x}{2\sqrt{Dt}}}^{\infty} \phi\left(t - \frac{x^2}{4D\mu^2}\right)e^{-\mu^2}d\mu$$

where

$$\mu = \frac{x}{2\sqrt{D(t-\lambda)}}$$

In this case

$$C(x,t) = \frac{2}{\sqrt{\pi}}\int_{\frac{x}{2\sqrt{Dt}}}^{\infty} L_0\left(1 - \cos\left(\frac{2\pi}{T}\left(t - \frac{x^2}{4D\mu^2}\right)\right)\right)e^{-\mu^2}d\mu$$

$$= L_0\left[1 - \mathrm{erf}\left(\frac{x}{2\sqrt{Dt}}\right) - \frac{2}{\sqrt{\pi}}\int_{\frac{x}{2\sqrt{Dt}}}^{\infty} \cos\left(\frac{2\pi}{T}\left(t - \frac{x^2}{4D\mu^2}\right)\right) e^{-\mu^2} d\mu\right]$$

Since by a known definite integral

$$\frac{2}{\sqrt{\pi}}\int_0^{\infty} \cos\left(\frac{2\pi}{T}\left(t - \frac{x^2}{4D\mu^2}\right)\right) e^{-\mu^2} d\mu = \cos\left(\frac{2\pi}{T}t - \sqrt{\frac{\pi}{DT}}x\right) e^{-\sqrt{\frac{\pi}{DT}}x}$$

Then

$$C(x,t) = L_0\left[1 - \cos\left(\frac{2\pi t}{T} - \sqrt{\frac{\pi}{DT}}x\right)\exp\left(-\sqrt{\frac{\pi}{DT}}x\right)\right]$$

$$- L_0\left[\mathrm{erf}\left(\frac{x}{2\sqrt{Dt}}\right) + \frac{2}{\sqrt{\pi}}\int_0^{\frac{x}{2\sqrt{Dt}}} \cos\left(\frac{2\pi}{T}\left(t - \frac{x^2}{4D\mu^2}\right)\right) e^{-\mu^2} d\mu\right]$$

The second term is a transient disturbance caused by starting the oscillations of source at $t$=0, and it dies away as $t$ increases. In the steady state, for a finite value of $x$,

$$C(x,t) = L_0\left[1 - \cos\left(\frac{2\pi t}{T} - \sqrt{\frac{\pi}{DT}}x\right)\exp\left(-\sqrt{\frac{\pi}{DT}}x\right)\right].$$

**Supplemental references:**


1 Carslaw, H. & Jaeger, J. Conduction of heat in solids. *Oxford: Clarendon Press* (1947).

2 Hu, B. & Tu, Y. Precision sensing by two opposing gradient sensors: how does Escherichia coli find its preferred pH level? *Biophys J* **105**, 276-285 (2013).

3 Hu, B. & Tu, Y. Behaviors and strategies of bacterial navigation in chemical and nonchemical gradients. *PLoS Comput Biol* **10**, e1003672 (2014).

**Movie S1.** An example video of wild-type *E. coli* HCB1 cells swimming up the potassium gradient in a microfludic device. The left side is the source channel.

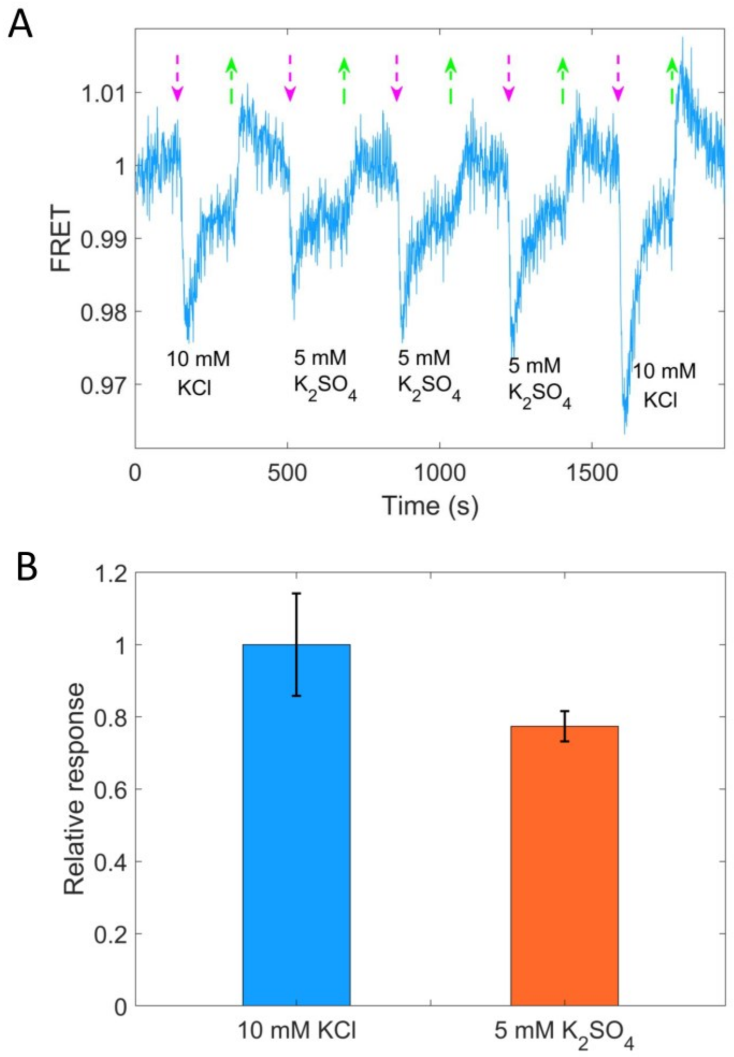


Fig. S1. **A.** The chemotactic response of the wild-type strain (HCB1288-pVS88) to 10 mM KCl and 5 mM $K_2SO_4$. The vertical purple (green) arrows denote the moment of adding (removing) stimulus. **B.** Quantitative comparison among the responses to 10 mM KCl and 5 mM $K_2SO_4$. The errors denote SEM.

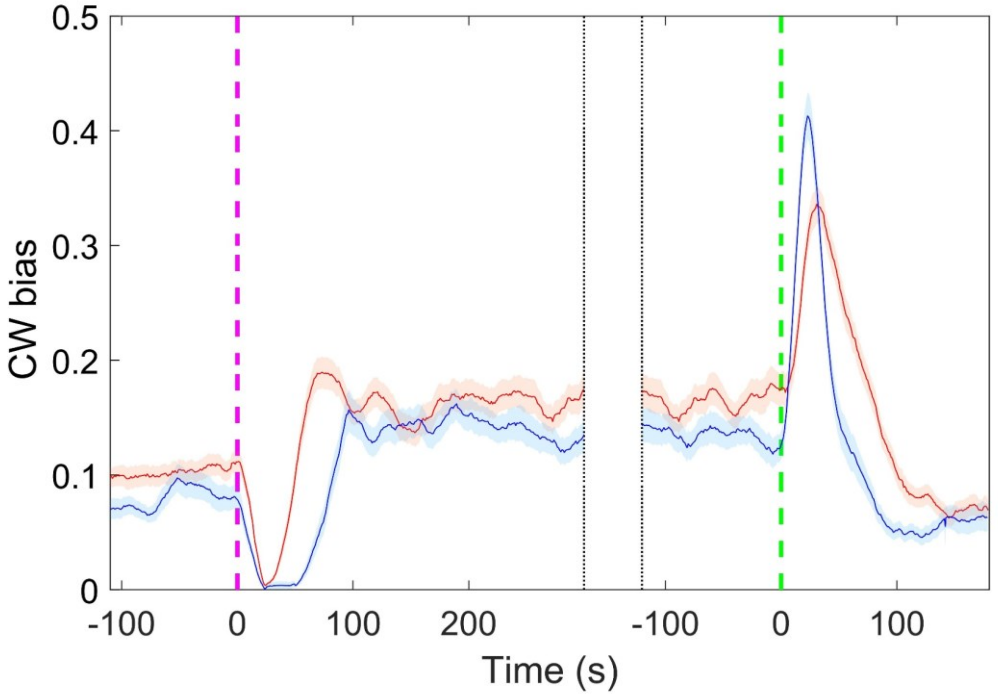


Fig. S2. The response of motor CW bias to 15 mM $K_2SO_4$ (red line) and 30 mM KCl (blue line) for the wild-type strain. There are 83 motors from 5 samples to 30 mM KCl and 91 motors from 4 samples to 15 mM $K_2SO_4$. The shaded areas denote SEM.

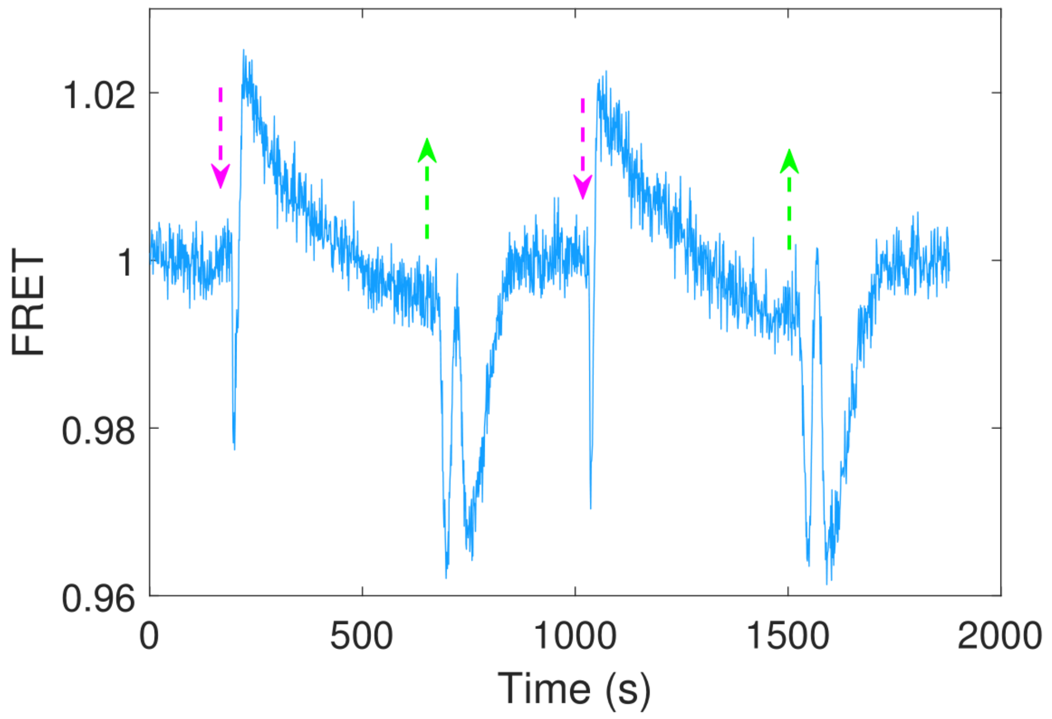


Fig. S3. The chemotactic response of the Tar-only strain (HCB1414-pLC113-pVS88) to 40 mM sodium benzoate at pH=7.0. The vertical purple (green) arrows denote the moment of adding (removing) stimulus. The response to the removal of sodium benzoate seems to be a superposition of an attractant and a repellent response, the reason for which deserves to be further explored.

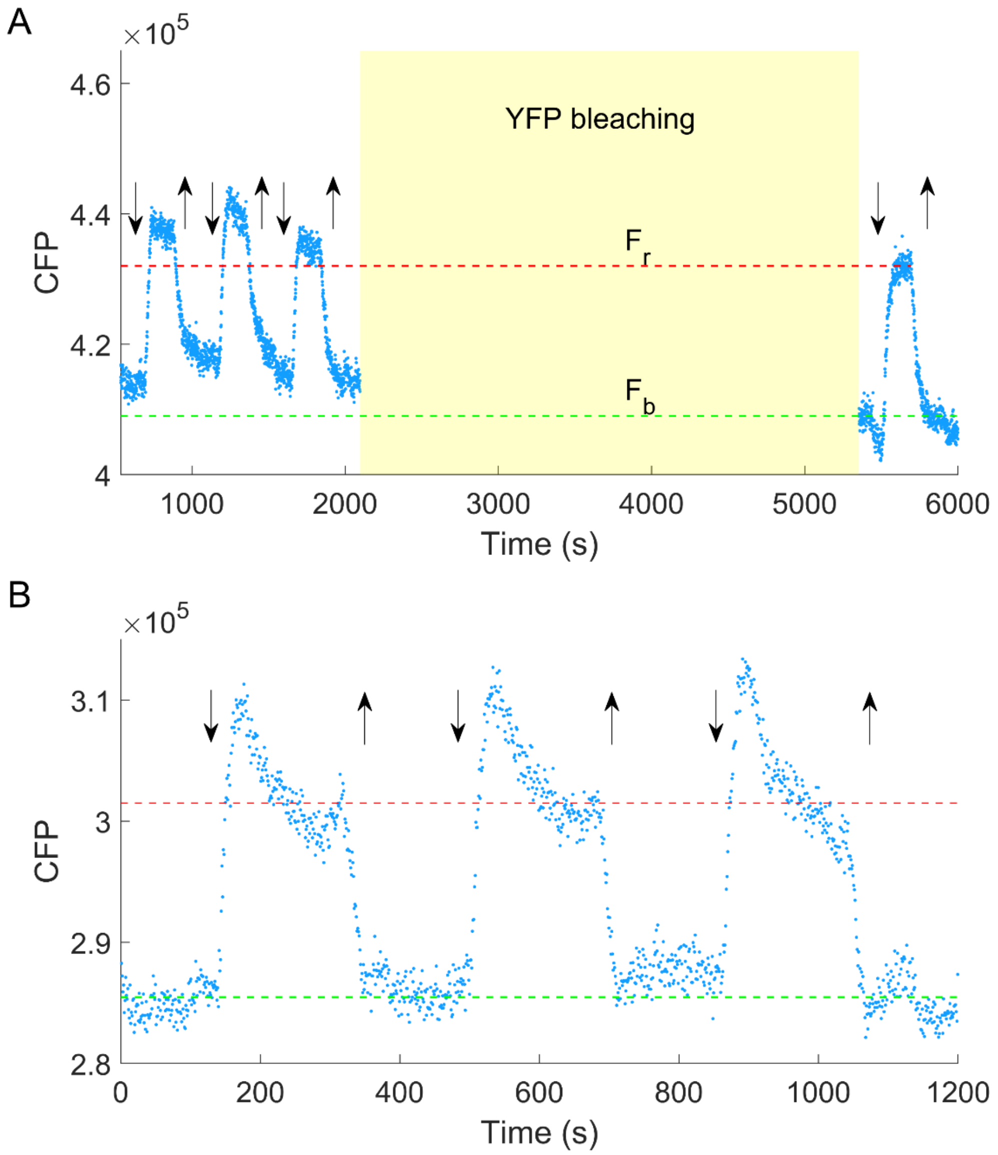


Fig. S4. The CFP intensity response to 30 mM KCl. **A.** The CFP intensity response of HCB1414-pVS88 to 30 mM KCl before and after YFP bleaching (50 min, yellow shaded area). Blue dots are experimental data. $F_r$ and $F_b$ are the CFP intensities for cells in motility medium (green dashed line) and 30 mM KCl (red dashed line), respectively. **B.** The CFP intensity response of HCB1288-pVS88 to 30 mM KCl. Blue dots are experimental data. The CFP intensity for cells in motility medium is denoted with the green dashed line. The theoretical fluorescence enhancement (red dashed line) was calculated by multiplying the level of the green dashed line by $F_r/F_b$ in A. The vertical purple (green) arrows denote the moment of adding (removing) 30 mM KCl. The background measured with HCB1288-pVS18 was subtracted.

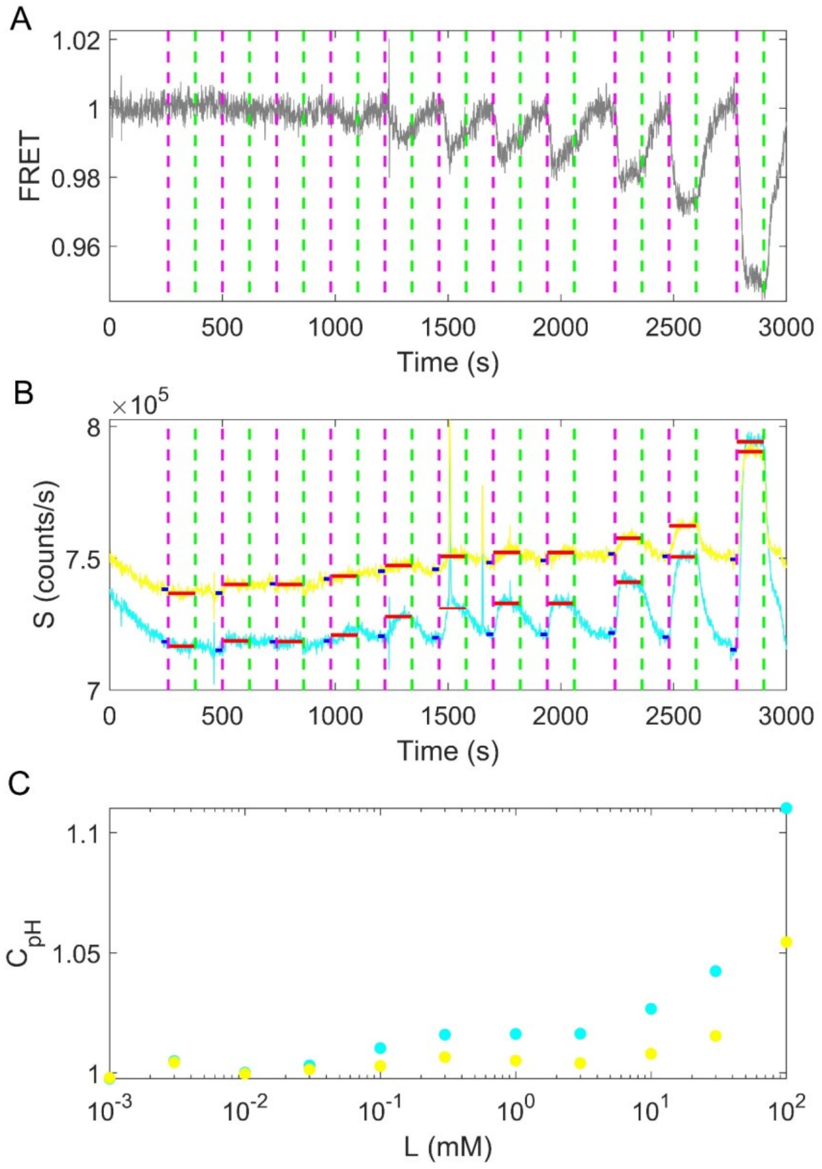


Fig. S5. The response of the no-receptor mutant (HCB1414-pVS88) to different concentrations of KCl. **A.** The original FRET signal of the no-receptor mutant in response to KCl. The vertical purple (green) dashed lines indicate the moment of adding (removing) KCl in the following order: 0.001 mM, 0.003 mM, 0.01 mM, 0.03 mM, 0.1 mM, 0.3 mM, 1 mM, 3 mM, 10 mM, 30 mM and 100 mM. **B.** The original CFP (cyan solid line) and YFP (yellow solid line) signals of the no-receptor mutant in response to KCl. The vertical purple (green) dashed lines represent the same events as in (**A**). The horizontal blue (red) solid lines represent the PMT signal values before (after) the addition of KCl at special concentrations. **C.** The relationship between the ratio of PMT signal post- to pre-KCl addition and the concentration of KCl for CFP (cyan dots) and YFP

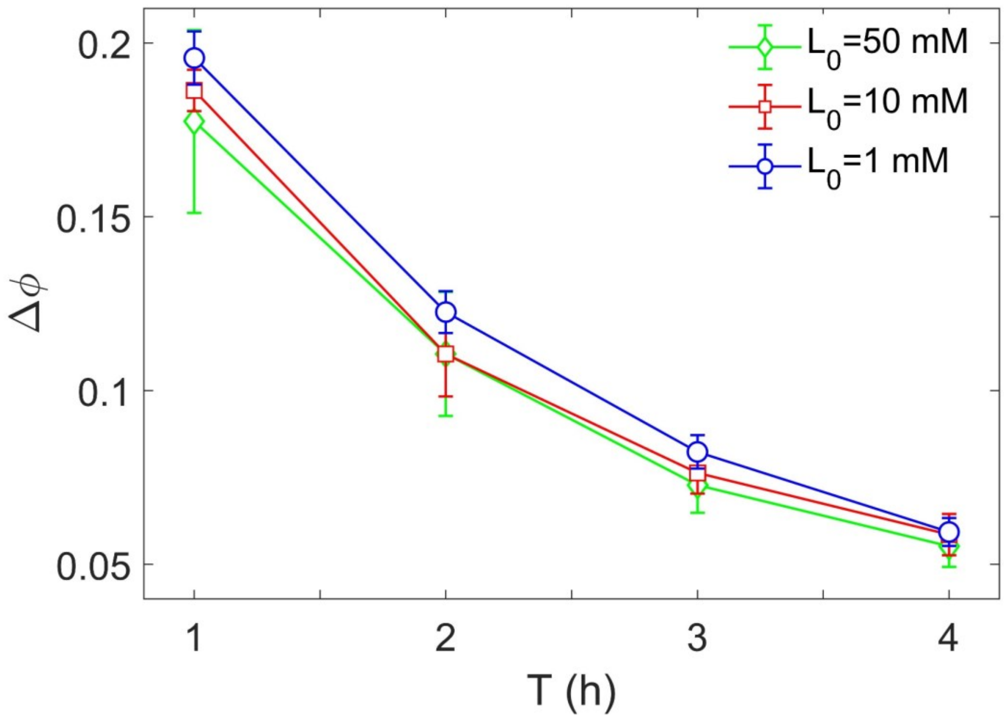


Fig. S6. The relation between phase delay ($\Delta\phi$) and driving periods ($T$) with different $L_0$.

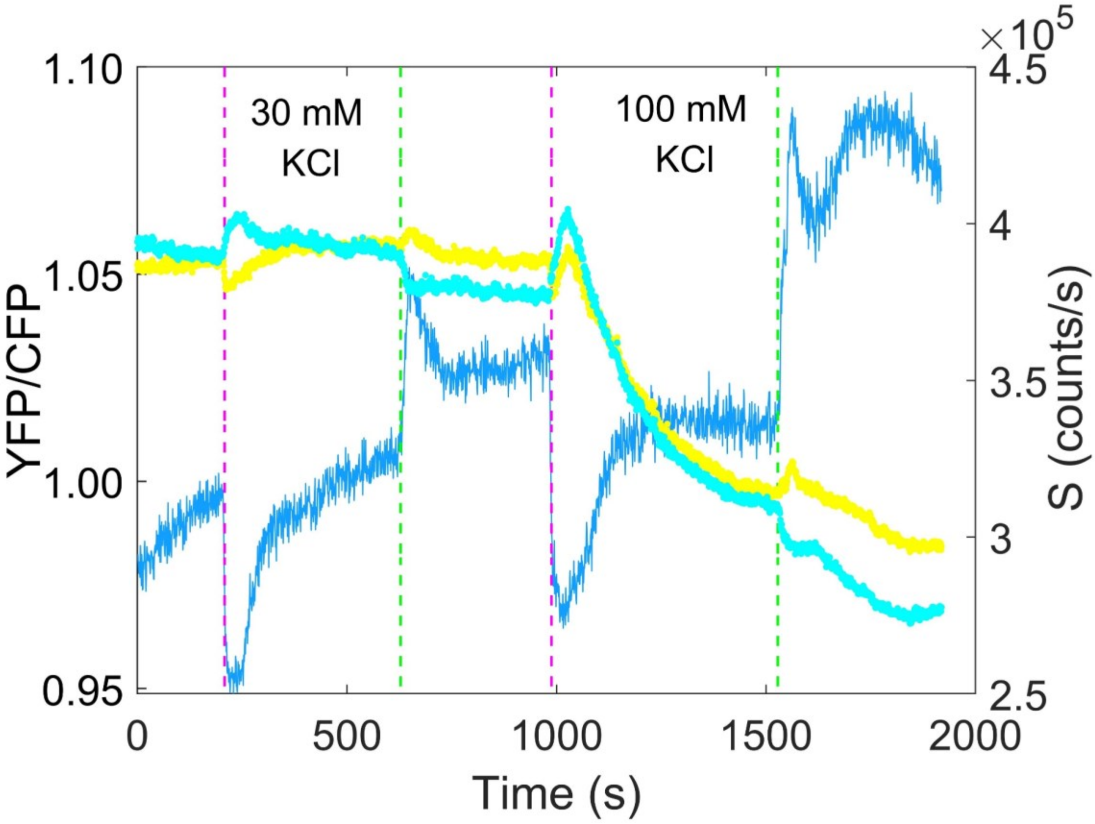


Fig. S7. The response of wild-type strain (HCB1288-pVS88) to KCl. The blue solid line denotes the ratio of YFP to CFP, while the cyan and yellow dots represent the PMT signals from CFP and YFP channel respectively. The vertical purple (green) dashed lines indicate the moment of adding (removing) 30 mM and 100 mM KCl in order.

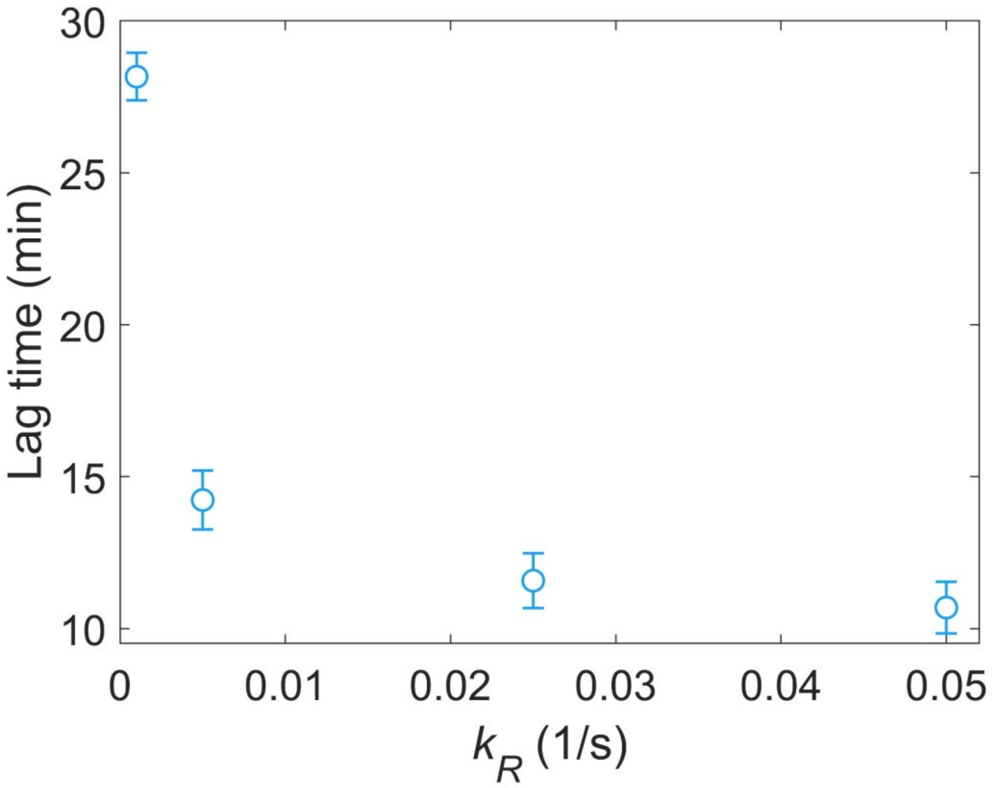


Fig. S8. The relationship between the lag time ($\Delta\phi \cdot T$) and the methylation rate $k_R$.
